# Epigenome-based Splicing Prediction using a Recurrent Neural Network

**DOI:** 10.1101/2020.02.03.932251

**Authors:** Donghoon Lee, Jing Zhang, Jason Liu, Mark B Gerstein

## Abstract

Alternative RNA splicing provides an important means to expand metazoan transcriptome diversity. Contrary to what was accepted previously, splicing is now thought to predominantly take place during transcription. Motivated by emerging data showing the physical proximity of the spliceosome to Pol II, we surveyed the effect of epigenetic context on co-transcriptional splicing. In particular, we observed that splicing factors were not necessarily enriched at exon junctions and that most epigenetic signatures had a distinctly asymmetric profile around known splice sites. Given this, we tried to build an interpretable model that mimics the physical layout of splicing regulation where the chromatin context progressively changes as the Pol II moves along the guide DNA. We used a recurrent-neural-network architecture to predict the inclusion of a spliced exon based on adjacent epigenetic signals, and we showed that distinct spatio-temporal features of these signals were key determinants of model outcome, in addition to the actual nucleotide sequence of the guide DNA strand. After the model had been trained and tested (with >80% precision-recall curve metric), we explored the derived weights of the latent factors, finding they highlight the importance of the asymmetric time-direction of chromatin context during transcription.

**Author Summary:** In humans, only about 2% of the genome is comprised of so-called coding regions and can give rise to protein products. However, the human transcriptome is much more diverse than the number of genes found in these coding regions. Each gene can give rise to multiple transcripts through a process during transcription called alternative splicing. There is a limited understanding of the regulation of splicing and the underlying splicing code that determines cell-type-specific splicing. Here, we studied epigenetic features that characterize splicing regulation in humans using a recurrent neural network model. Unlike feedforward neural networks, this method contains an internal memory state that learns from spatiotemporal patterns – like the context in language – from a sequence of genomic and epigenetic information, making it better suited for characterizing splicing. We demonstrated that our method improves the prediction of spicing outcomes compared to previous methods. Furthermore, we applied our method to 49 cell types in ENCODE to investigate splicing regulation and found that not only spatial but also temporal epigenomic context can influence splicing regulation during transcription.

## Introduction

Alternative splicing of pre-messenger RNA plays an integral role in diversifying the transcriptome. This process is more pervasive in higher eukaryotes and is estimated to affect approximately 95% of protein-coding genes in humans [1,2]. Accurate characterization of the process by which multiple functional protein products are produced from a single gene is crucial for understanding the function of the transcriptome [3].

Recent discoveries have revealed that splicing occurs predominantly during transcription in humans [4–8]. Nascent RNA is almost immediately spliced upon transcription [9,10] and introns are mostly spliced out during transcript elongation. This timing suggests that the recruitment of splicing factors and spliceosome assembly, detection of exon-intron boundaries, and modulation of alternative splicing must occur at the same time scale as transcription [9].

Co-transcriptional splicing indicates a key observation that splicing takes place progressively in the direction of RNA transcription, rather than processed simultaneously after transcription. As a result, the contexts of guide DNA, nascent RNA, and its immediate folded structure progressively change as RNA polymerase II (Pol II) moves along the guide DNA strand [11] and may influence splicing regulation. Furthermore, co-transcriptional splicing signifies the physical proximity of the spliceosome assembly to Pol II and other transcriptional machinery [9]. Pol II physically interacts with nucleosomes and its histone modifications around them, modulating the transcription rate [12].

DNA sequence alone may not contain sufficient information to process alternative splicing deterministically [13]. Djebali et al. [4] and many others have shown that there is an enrichment of chromatin marks around spliced exons, suggesting the role of epigenetic modifications during context-dependent modulation of alternative splicing [14,15]. For example, exonic boundaries are characterized by increased levels of nucleosome density and positioning [16–18], DNA methylation [19,20], and strong enrichment of specific histone modifications including H3K36me3, H3K79me1, H2BK5me1, H3K27me1, H3K27me2, and H3K27me3 [16,17,21–23]. In addition, a recent genome-wide survey of alternative splicing showed that DNA methylation can either enhance or silence exon recognition in a context-dependent manner [24]. Furthermore, studies have shown that there is significant regulatory crosstalk between histone modifications during transcriptional elongation [12].

Despite many efforts to characterize the splicing regulatory code both experimentally and computationally, we have yet to understand how the cell type-specific epigenomic context is utilized during co-transcriptional splicing. Previous computational methods on splicing have largely focused on discovering novel splice junctions based on RNA sequencing (RNA-seq) alignments [25,26], utilizing machine learning approaches [27,28] including deep neural networks [29]. Only a limited set of tools can model splicing regulation based on genomic sequences and select RNA features [30–32]. Moreover, studies on splicing regulation have focused heavily on identifying mutations that land within splice sites (SSs), cis-acting splicing regulatory elements, and trans-acting splicing factors [30,33]. The extent, nature, and effects of the epigenetic context in splicing regulation remain unsolved.

In this study, we propose a new computational approach to characterize the role of epigenetic modifications during co-transcriptional splicing. To build an interpretable model, we adopted a recurrent neural network (RNN) architecture, which to some degree resembles the physical characteristics of co-transcriptional splicing (Figure 1). The model can learn from a temporal sequence of epigenetic contexts, similar to how epigenetic contexts progressively change as Pol II moves forward along the guide DNA strand during co-transcriptional splicing. The RNN model allows us to predict the inclusion of exons based on adjacent DNA sequences and epigenetic modifications. Moreover, the physical resemblance of the model allows us to interpret the trained model weight parameters and explore the spatio-temporal links between the guide DNA elements and the surrounding epigenetic modifications. In summary, we leveraged the mechanistic properties of co-transcriptional splicing to build an interpretable splicing model, and we explored the trained model to understand the underlying characteristics of the epigenetic context during co-transcriptional splicing.

**Figure 1.**
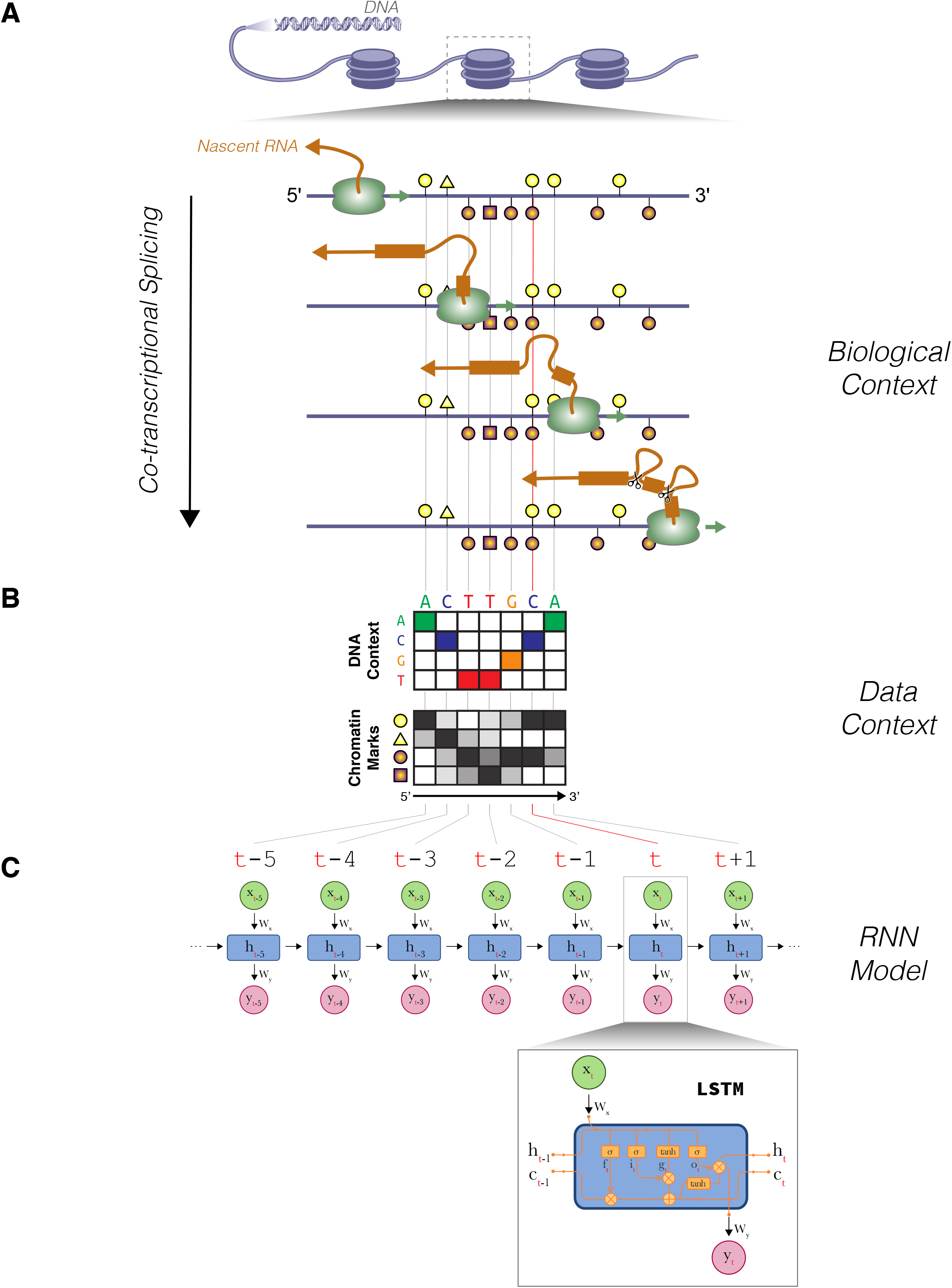
Overview of the co-transcriptional splicing model. Depiction of co-transcriptional splicing in terms of **(A)** biological context, **(B)** genomic and epigenomic data context, and how it relates to the **(C)** RNN model.

## Results

We first explore the epigenetic data context around known splice sites in depth. We then describe the model and rationale for applying the specific architecture. Finally, we use the model to further examine the effect of epigenetic context during co-transcriptional splicing.

### Distinct epigenomic signatures characterize splicing regulation

We studied the epigenetic context of alternative splicing by examining the enrichment of multiple histone modifications and DNA methylations around the exon-intron boundary. We mapped the epigenomic signatures around SSs of cassette exons at a base-pair resolution. We aggregated multiple histone modifications across 49 cell types in ENCODE and observed their enrichment as a function of distance from SSs (Figure 2A, B, Supplementary Figure 1, 2A, B). We found the most interesting trend within 100 bp of SSs for both the 3’ acceptor and 5’ donor. A strong enrichment pattern of H3K36me3 and H3K27me3 appeared around the exon boundary. Although studies have demonstrated a role for H3K36me3 in defining the exon-intron boundary [22,34], the dynamic interplay between other histone modifications has been overlooked. From the 3’ acceptor, peak enrichment occurred around 100 bp into the exon; at the 5’ donor, it was closer, at around 50 bp into the exon. We also observed a slight depletion of H3K27ac and H3K4me3 marks within 100 bp of the intron at the 3’ acceptor SS but not within the 5’ donor SS. Using Mann-Whitney-Wilcoxon tests, we confirmed that the relative elevation and depletion of epigenetic enrichment at the genomic segment containing the branching site (segment C) compared to the surrounding exons (Figure 2B, Supplementary Figure 2A, B). As this region contains a branch site, these histone marks may indicate a role in defining the branch point.

**Figure 2.**
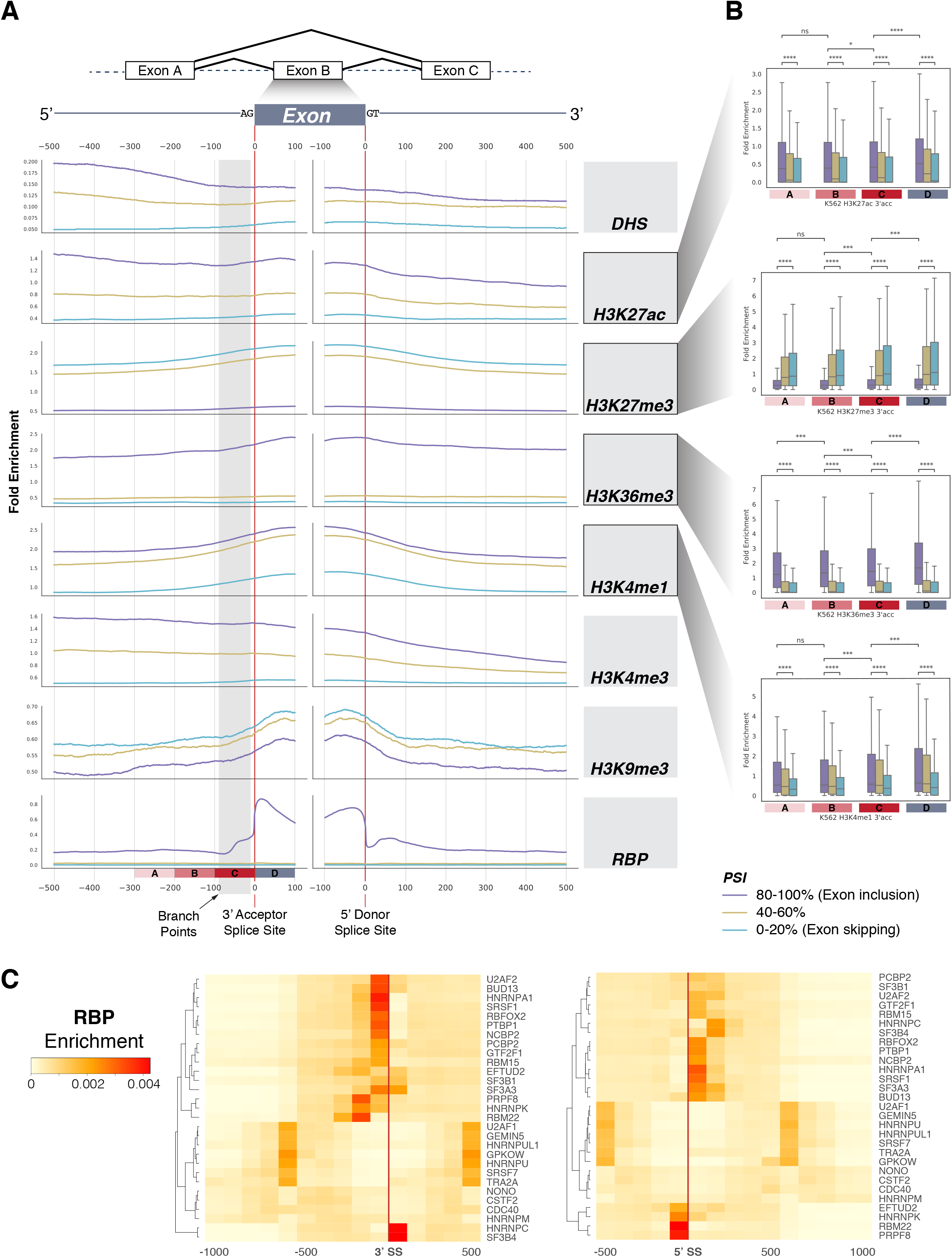
**(A)** Enrichment of various epigenomic marks of K562 at the exon-intron boundary. We aggregated histone modifications up to 500 bp upstream and downstream of intronic and exonic regions flanking 3’ and 5’ SSs for cassette exons across ENCODE cell types. High PSI indicates exon inclusion, mid PSI indicates exons with 40-60% PSI, and low PSI indicates exon skipping. **(B)** Statistical significance testing of epigenetic mark enrichment. Average histone modification enrichment at four exonic segments were compared based on PSI values. Mann-Whitney-Wilcoxon two-sided test, ns: 0.05 < p <= 1; *: 0.01 < p <= 0.05; **: 0.001 < p <= 0.01; ***: 0.0001 < p <= 0.001; ****: p <= 0.0001. **(C)** RBP enrichment across the exon-intron boundary.

### Enrichment of RNA-binding factors around splice sites

Alternative splicing regulation is an elaborate process that requires precise coordination of multiple splicing factors and enzymes. Studies have shown that RNA-binding proteins (RBPs) facilitate splicing regulation during transcription [35]. For example, the serine/arginine-rich splicing factor family member SRSF7 binds to poised exons and promotes the inclusion rate [36][37]. Another member of the serine/arginine-rich splicing factor family, U2AF1, is responsible for mediating the binding of U2 small nuclear ribonucleoprotein to the pre-mRNA branch site [38]. The recent release of the ENCODE project included enhanced CLIP experiments (eCLIP) datasets that span 112 RBPs from K562 and HepG2 cell types. As sequence-specific RBPs have been shown to facilitate splicing regulation in a context-specific manner [15], we investigated their spatial relationship to both the 5’ donor and 3’ acceptor splicing sites. Specifically, we investigated the enrichment of splicing factors (n=29) and their relative distance to these sites. We observed that, on average, splicing factors show preferential binding to the intronic side of the splicing site in both 3’ acceptor and 5’ donor SSs (Supplementary Figure 2C). Furthermore, we found that splicing factors may show slightly different patterns in their spatial binding preferences. In particular, hnRNP A1 and SRSF1 were enriched in the intronic region outside 3’ SSs whereas SF3B4 and hnRNP C were enriched in the exonic region (Figure 2C). At 5’ SSs, RBM22 and PRPF8 were bound at the exonic end, which has been shown to be critical for splicesome assembly [39,40].

### Correlating epigenomic signatures to exonic expression

We tested whether histone modifications have any effect on inclusion and expression of alternative exons. We observed a trend where enrichment of H3K36me3 at the exon-intron boundary was positively correlated with exonic expression, whereas H3K27me3 marks showed the opposite trend (Figure 3A, B, Supplementary Figure 3). Compared to excluded or nominally expressed alternative exons, highly expressed spliced exons had statistically significant enrichment of H3K36me3 and depletion of H3K27me3 at their exon-intron boundary (Figure 3C). The contrasting trend and the correlation of these histone methylations to exonic expression suggest that the splicing code may be directly or indirectly encoded within the epigenomic context.

**Figure 3.**
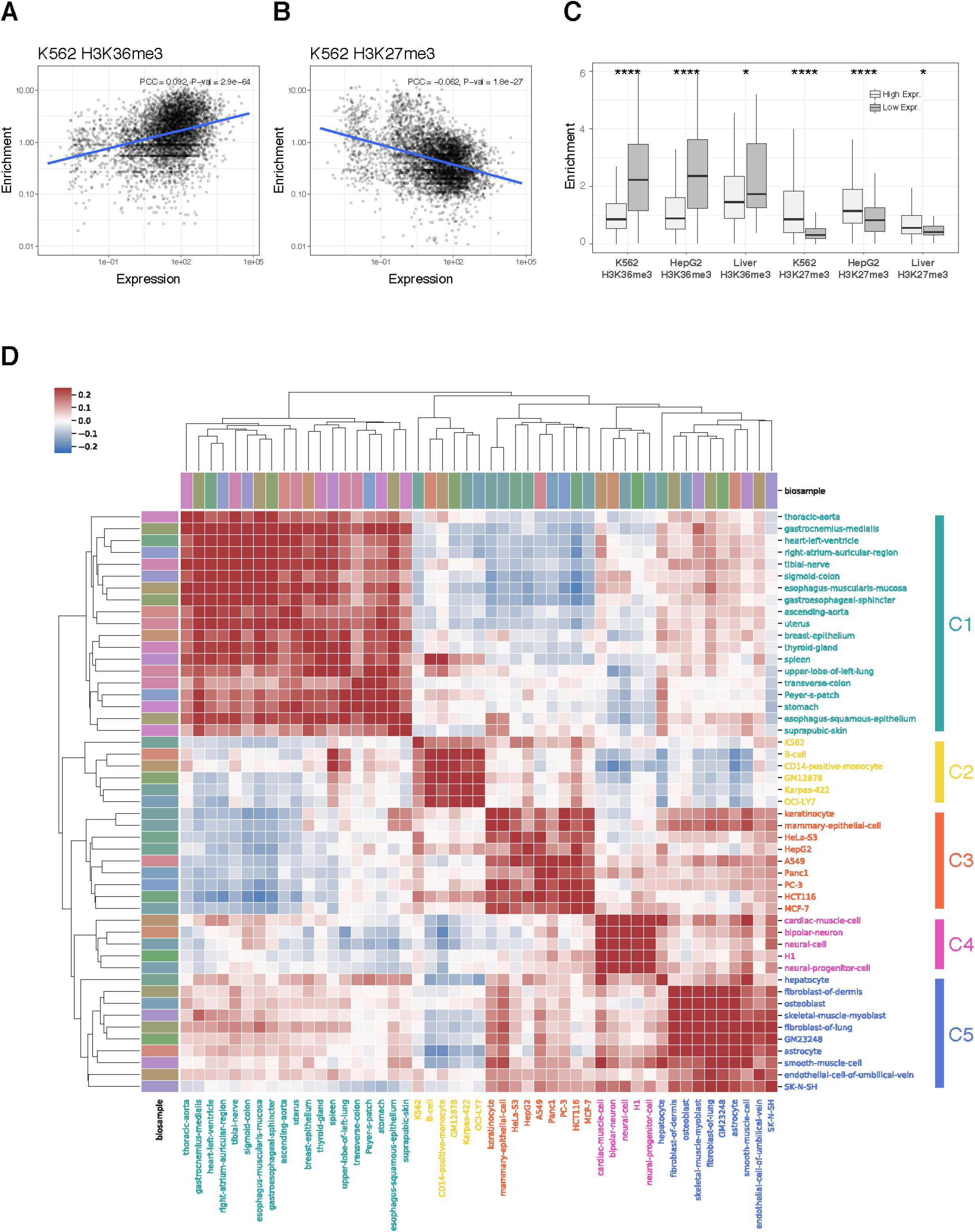
Correlation of exonic expression to **(A)** H3K36me3 and **(B)** H3K27me3. The line represents a linear regression model fit, and the shaded band represents 95% confidence interval. **(C)** Alternative exons were grouped by expression level and their relative histone enrichment was compared near the SSs. Asterisks represents statistical significance using the Wilcoxon rank sum test; (*) P <= 0.05, (**) P <= 0.01, (***) P <= 0.001, (****) P <= 0.0001. **(D)** Hierarchical clustering of similarity based on PSI across 49 ENCODE biosamples. The results are clustered into five categories of cell types.

### Clustering biosamples based on splicing patterns

Previous studies have shown that various epigenomic marks are correlated across similar tissues and cell types [41]. It is now widely accepted that the transcriptional regulatory circuitry of a particular cell type is reflected in its epigenetic landscape. To explore the potential linkage between epigenetic regulation and tissue-specific splicing, we examined splicing patterns across 49 ENCODE biosamples. Based on a similarity of percent-splice-in (PSI) values for all coding exons (n=185,405), we clustered biosamples into five categories using hierarchical clustering (Figure 3D). Splicing patterns were highly correlated among tissue types from the same cell-of-origin, reproducing similar clustering results based on epigenetic marks. For example, blood-lineage cell types formed cluster C2 whereas brain and neural cells were clustered in cluster C4. Moreover, we observed that cancerous cell lines cluster together in cluster C3.

In addition to using the PSI similarity matrix to cluster cell types into categories, we can project the cells onto a low-dimensional cell space using principal component analysis (PCA). We measured alternative splicing patterns in terms of exonic expression level (fragment per kilobase per million reads mapped, FPKM) across diverse ENCODE cell types and examined how cells are placed in the context of others. Interestingly, we observed that cancer-related cell lines were located proximal to each other in the PCA cell space (Supplementary Figure 4).

### Modeling splicing regulation: key characteristics of an RNN architecture

To investigate the latent representation of splicing instruction encoded within the epigenomic context, we aimed to construct a predictive model of splicing. We opted for an RNN architecture, which has proven successful in various sequential information processing and prediction tasks such as natural language processing and translation [42–44], to explore the contribution of the epigenomic context to the regulation of alternative splicing.

We start by describing a simple RNN, which shares many of the features we intend to model. A simple RNN is made of many recurrent neurons that are sequentially linked to each other. A neuron at specific time point *t* is influenced by previous time point *t*−1, combining some relationship of the current input *x*_*t*_ with the previous hidden state *h*_*t*−1_.

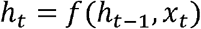

where *h*_*t*_ is hidden state at time *t* and *x*_*t*_ is input variable at time *t*. If we suppose the activation function as a hyperbolic tangent for a simple RNN, the state at time *t* can be represented as

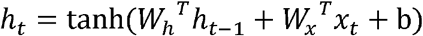

where *W*_*h*_ and *W*_*x*_ are the weight of the hidden state and input variable, respectively, and b is the bias vector. The output can be expressed in terms of an output weight matrix, *W*_*y*_, and a hidden state at time *t*, *h*_*t*_:

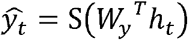

where *S* is sigmoid function:

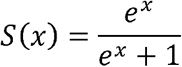

This time-dependency allows us to explore the complex contextual relationship between features. In particular, we adopted the long short-term memory (LSTM) [45] model to describe an RNN architecture. In principal, a simple RNN allows us to model a time-dependent task from sequential data. However, in practice, the simple model suffers from the problem of vanishing gradients, where the gradients responsible for updating weights with respect to the partial derivative of error function becomes negligible in a long sequence and hampers the model from learning long-term time dependencies. Therefore, we used both LSTM and gated recurrent unit (GRU), which have many of the same simple intuitive properties of the simple RNN but allow learning from longer sequences. The LSTM is an extension of the same idea that includes more sophisticated gates, which allows the cell to retain long-term memory between cells while avoiding the problem of vanishing gradients when training the network. The specific equations for the LSTM model we adopted is shown in the Methods.

### Modeling splicing regulation: How the RNN architecture fits the problem

The rationale for applying an RNN to our model is that (1) an RNN is optimized for processing sequential information like genomic sequences and epigenomic profiles along genomic coordinates, (2) an RNN has a time-direction resembling how RNA is transcribed by RNA polymerase in the 5’ to 3’ direction, (3) temporal memory cells of an RNN allow the model to learn about complex context-dependent relationships among epigenomic features, such as the influence of features and input seen at t-1 on the neural cell at time t, and (4) an RNN is very flexible with the type of input and output data and therefore can easily integrate heterogeneous sequential information. Not surprisingly, researchers recently have applied RNN models to the area of genomics to predict non-coding DNA function [46] and to detect exon junctions [47]. Moreover, since the mechanics of the RNN calculation is somewhat parallel to the actual spatial and temporal dependency found in co-transcriptional splicing, the overall results from the trained model are more readily interpretable. The data processing and implementation of the predictive models are collected in a package named <u>E</u>pigenome-based <u>S</u>plicing <u>P</u>rediction using <u>R</u>ecurrent <u>N</u>eural <u>N</u>etwork (ESPRNN; available at https://github.com/gersteinlab/esprnn). Using our method, we attempted to decipher context-dependent effects of various epigenomic features on splicing for both canonical (e.g., dinucleotide GT for 5’ donors and AG for 3’ acceptors) and non-canonical SSs. Our model is especially useful since splicing signals are not only enriched at the splice site but often found up and downstream of splice sites.

### Modeling splicing regulation: Initial evaluation

We used ESPRNN to predict alternate usages of cassette exons (inclusion or exclusion of exons), the most common form of alternative splicing events [48], using DNA sequences and epigenomic signals adjacent to SSs (Figure 4A). We used the exon definition of splicing, which is considered to be the dominant mechanism in higher eukaryotes [49]. Our model had an average F1 score (harmonic mean of the precision and recall) of 0.8472 for the LSTM-based model across cell types [0.8757 for the GRU-based model] using five core histone modification tracks (Figure 4B). The average F1 score marginally increased to 0.8573 when using 17 histone, chromatin accessibility, DNA methylation, and nucleosome density profiles.

**Figure 4.**
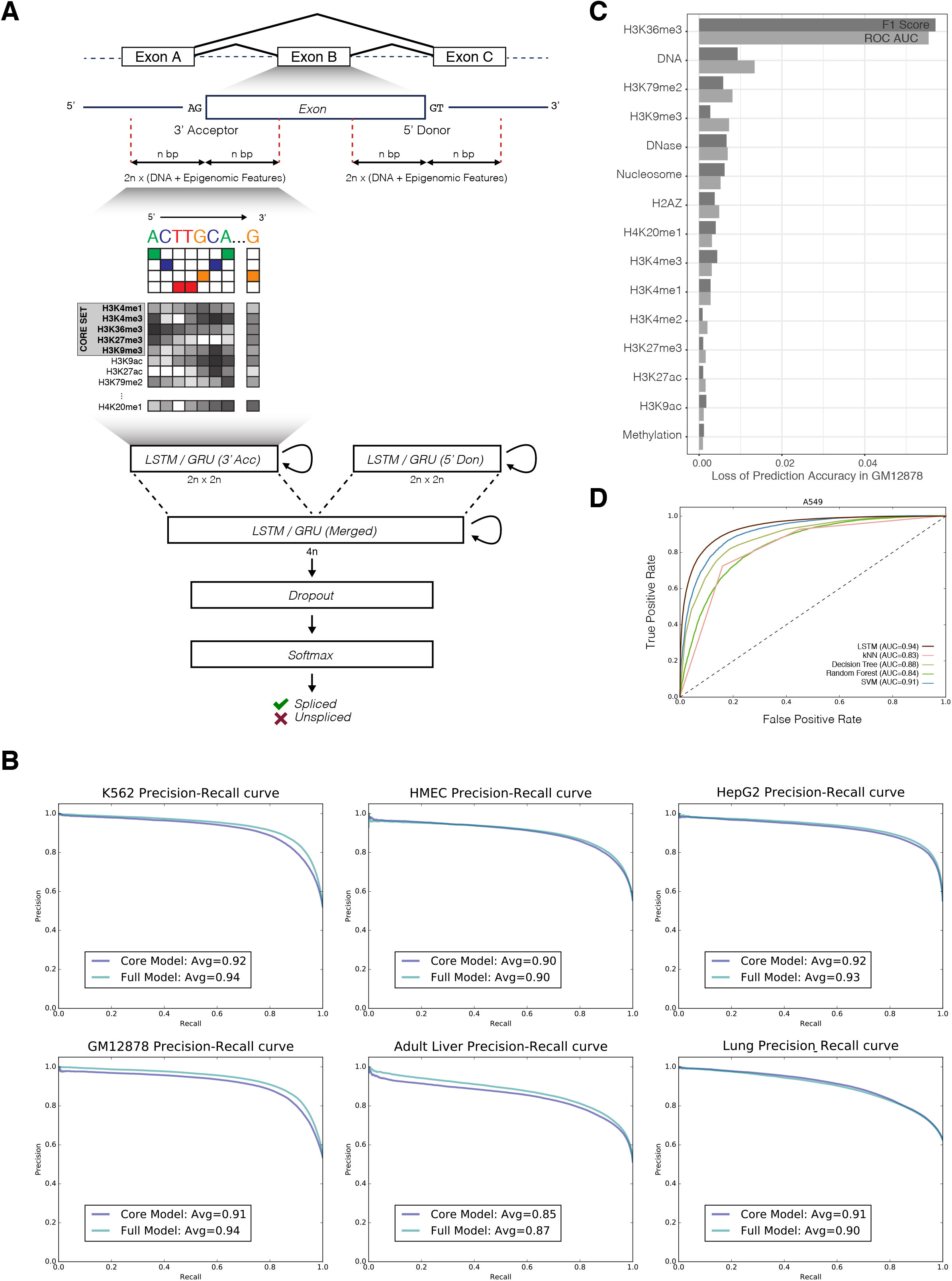
**(A)** Overview of the ESPRNN model. The model is composed of two recurrent layers. Inputs from 3’ and 5’ SSs are separately processed in the first recurrent layer and then merged in the next recurrent layer. A softmax classifier is used to determine the inclusion of the exon. Using genomic sequences and epigenomic contexts as input, the alternative usage of the exon is predicted. **(B)** Precision-recall curves from six different ENCODE cell types. **(C)** Epigenetic features that contribute to splicing regulation. The order and magnitude of importance was determined using leave-one-out analysis and loss of the ROC AUC was calculated when training the model lacking a particular feature. **(D)** Comparison of LSTM model with other models based on k-nearest neighbor, support vector machine, decision tree, and random forest algorithms.

We performed the splicing prediction with or without the RBP profile and measured how much predictive performance is gained from additional information. We observed a marginal improvement in predictive performance when RBP binding profiles were added to the baseline model (measured in improvement of F1 score from 0.84 to 0.86) (Supplementary Figure 9A, B). This suggests RBP binding information may be redundant and already represented in the epigenetic features. We also compared prediction results from normal cell types to those from cancerous cell lines. Since previous studies on cancer-specific alternative splicing [50,51] have suggested potential linkage of aberrant splicing events to the disease risk [52–55], we expected to see differences in splicing regulation between normal and cancerous cell types. However, we did not observe a significant difference in prediction performance between normal and cancerous cell types (average F1 score for normal biosamples: 0.8465, cancerous biosamples: 0.8765). We also cross-tested a model trained from one cell type to another. After we fit our model to one cell type, we transferred the fitted weights and model parameters to predict splicing on other cell types. When we tested between cell types from the same cell-of-origin (e.g., train on adult liver model and test on HepG2 data, train on lung model and test on A549 data), we did not observe a significant difference in predictive performance. However, we observed a moderate reduction in splicing prediction performance when we cross-tested cells from different cell-of-origin (Supplementary Figure 5B, F1 score is better metric for comparing cross-cell testing due to class imbalance across cell types). Thus, the epigenomic regulatory landscape around SSs appears to be generally conserved across cell types. Moreover, we compared the classification performance to other models based on random forest and k-nearest neighbors and found that our model was superior in terms of classification accuracy (Figure 4D, Supplementary Figure 7).

We tried to measure the contribution of each individual epigenetic feature to splicing in a number of ways. (1) We performed an empirical analysis via a leave-one-out strategy. Using GM12878 as an example, we first built a reference model based on all available epigenetic features. By removing one variable at a time, we then measured the mean decrease in F1 score and area under the receiver operating characteristic curve (ROC AUC), as an indicator of variable importance (Figure 4C). (2) Alternatively, we trained a DNA-only model using DNA sequence features only and compared to a “baseline model.” The baseline model was trained using DNA sequence features plus additional chromatin accessibility (DHS) and 6 histone marks. Here, we observed a significant loss of predictive performance in the DNA-only model (13% reduction in F1 score) (Supplementary Figure 6A). (3) Next, starting from the DNA-only model, we added one epigenetic feature at a time to measure the information gain from each feature (Supplementary Figure 6B). While the addition of some epigenetic features like H3K27ac increased the variability in prediction performance, an active mark H3K36me3 or a repressive mark H3K27me3 was the most informative at predicting splicing. Moreover, the combination of both H3K36me3 and H3K27ac further improved the prediction performance compared to other pairs (Supplementary Figure 6C). We observed that the combination of H3K36me3 and H3K27ac features together contributed more than when they were used individually (Supplementary Figure 6D).

Overall, we found H3K36me3 to be the most important variable in predicting splicing. This observation coincides with previous studies reporting that H3K36me3 recruits the splicing factors PTB [34] and SRSF1 [56] to facilitate splicing. Interestingly, one of the top predictors of splicing was H3K79me2, which was previously shown to associate with H3K36me3 at gene bodies [57]. H3K9me3, a histone modification that can recruit adaptor proteins like HP1 to facilitate splicing factors [24], was also ranked among the top predictors.

### Interpretation of weights of the splicing model

Since the model follows the physical layout of splicing regulation, one can examine the trained model and learn from the trained weights how each epigenetic feature contributes to splicing regulation. To interpret the splicing model, we designed an LSTM-based model composed of only one hidden state and trained for a longer period (400 epochs). We made sure that this simplified model performs nearly as well at predicting splicing as our main model (usually after >20 epochs of training, Supplementary Figure 8A). We also made sure that the overall predictive performance of the simplified model is stable after approximately 100 epochs (Supplementary Figure 8B, C). When we analyzed the simplified model, we found that the trained weights of various gates at the recurrent unit showed that open chromatin (DHS), H3K27ac, K3K36me3, and H3K4me1 are weighted more highly than other epigenetic features -- as expected (Supplementary Figure 8D). We also noticed that H3K27me3 and K3K9me3 were negatively weighted at the input gate, suggesting that these features have a negative impact on exon inclusion, consistent with our previous findings.

### Influence of temporal epigenetic context on splicing regulation

We specifically designed our splicing model to represent the physical layout of splicing regulation, where a sequence of chromatin contexts is fed progressively to the model. Therefore, the model takes into account the temporal direction (progression from 5’ to 3’ in direction). To show that model has learned this asymmetric temporal relationship of epigenetic features, we first trained a baseline model (in the normal 5’ to 3’ direction) and then fed a series of epigenetic signals in a “reverse” order (3’ to 5’ in direction) as input to it. We examined how the model prediction behaved in this context. If the model was agnostic to the temporal direction of features, both forward and reverse input features should give the same predictive power. By using a model based on a single histone feature, H3K36me3, we observed a moderate decrease in prediction performance upon reversal of the epigenetic feature (Supplementary Figure 9), with an F1 score decreasing from 0.78 to 0.77 and ROC AUC decreasing from 0.87 to 0.85. While we suspect there are some level of redundancy across different epigenetic marks and some marks are independent of their temporal direction, our results suggest the importance of temporal direction of epigenetic features in the context of splicing.

## Discussion

Our prediction model revealed that the epigenomic signature of an SS plays a large role in determining the splicing outcome. In addition, the positive results suggest that our model can be extended to predict the full transcriptomic composition from a genomic and epigenomic context. We expect that we could further improve the proposed model by adding more deep hidden layers and increasing the number of training samples by utilizing the full set of available epigenomic data in the ENCODE project. Our approach does contain some limitations, as it is still challenging to visualize and evaluate the multi-dimensional context of the weight matrix in the trained model. We could apply dimensionality reduction techniques to probe the latent representation of relationships between various epigenomic signals.

In this study, we used ENCODE polyA RNA-seq assays to measure splicing and exon-level expression; we note that this is an indirect measure of what is actually happening during transcription. RNAs are often unstable and may be subjected to many post-transcriptional modifications. RNA-seq measures the steady-state level of the transcript, accounting for both mRNA synthesis and decay. Future studies with a more direct measure of transcriptional rates, such as nuclear run-on assays like global run-on (GRO-seq) or bromouridine sequencing (Bru-seq), will allow us to accurately measure the effect of epigenomic context on splicing and, ultimately, on the transcriptional rate.

Future studies should focus on comparing splicing models from normal and cancer samples in the hope of illuminating the differences in the epigenomic landscapes of splicing regulation. Although splicing is an elaborate process, it could become pathogenic when misregulated [58,59]. Unsurprisingly, aberrant splicing events, which collectively referred to splicing events that could confer the risk of a disease, are often implicated in systemic diseases like cancer [51,60]. Aberrant splicing events based on mutations are relatively well characterized [54,60–62]; however, a large fraction of aberrant splicing events that have no direct mutational cause still remain unknown. Although our understanding of epigenomic context on splicing regulation is incomplete, our prediction model highlights that splicing is elaborately regulated via various epigenomic signatures. This suggests that epigenomic dysregulation may be closely linked to the onset of aberrant splicing. Thus, even though aberrantly spliced RNAs in healthy cells may be degraded by the mRNA surveillance system, epigenomic dysregulation may render this checkpoint system useless. Further studies on cell-type-specific and context-dependent splicing regulation will reveal whether epigenetic modulation can serve as a therapeutic method of complex disease in the future.

## Methods

### Dataset

The current release of the ENCODE dataset provides an unprecedented number of functional assays across broad biosample types, including primary cells and tissues. In this study, we leveraged both the breadth and depth of ENCODE, including assays for histone modification (chromatin immunoprecipitation sequencing, ChIP-seq), chromatin accessibility (DNase I hypersensitive sites sequencing, DNase-seq), RBPs (eCLIP), methylations (WGBS and RRBS) and gene expression (RNA-seq), to systematically probe the data-rich context of alternative splicing and its regulation. The list of accessions for experiments used in this study is found in Supplementary Table 1.

### Processing of RNA-seq data

To quantify levels of exon expression from RNA-seq data, we collected all raw sequencing reads from experiments tagged as reference epigenome series from the ENCODE portal. These reads were polyA plus long RNA-seq (200 bp or larger) from whole-cell fractions rather than nuclear or cytosolic fractions. To minimize potential batch effects and sample bias, we carefully selected untreated experiments from the reference epigenome series. As of November 2019, there are 81 cell and tissue types (covering 49 unique biosamples) in the reference epigenome series, including both RNA-seq and ChIP-seq of H3K4me1, H3K4me3, H3K36me3, H3K27ac, H3K27me3, and H3K9me3. We first aligned all RNA-seq data to the GRCh38 genome using RNA STAR (v 2.7.0). Since the model requires splice site annotation, we constructed exon annotation from GENCODE version 24 (to synchronize with ENCODE annotation) by extracting all unique exons with known protein-coding transcripts. We excluded exons that could ambiguously map to both chromosome X and Y. This analysis included 597,937 exons (185,405 unique exons after removing duplicates from isoforms) that averaged 28.01 exons per gene and 296.49 bp in length (150.92 bp in length for unique exons). We obtained read counts at each exon using HTSeq (v0.11.2) [63]. Based on read counts, we used a custom script (esprnn/preproc_calcExonFPKM.py) to calculate normalized exonic expression levels in FPKM. Our rationale for using the exonic expression was to intentionally make the model agnostic to the overall transcript level. Each exon was evaluated independently from other exons, and we counted the number of sequencing reads supporting the inclusion of a particular exon. The counts were normalized similar to how a gene’s expression is normalized by size of annotation and total number of mapped reads (FPKM). We binarized the exonic expression level (FPKM) using a threshold of one. Therefore, we only considered whether an exon has enough evidence supporting exon inclusion.

In addition to the exonic expression level, alternatively, we calculated a metric, PSI, to measure the level of splicing. PSI represents the fraction of the reads supporting exon inclusion from the split reads at the splice junction. We used a custom script (esprnn/scripts/calcPSI.sh) based on equations from Schafer et al. [64] to calculate PSI normalized by the size of read and exon annotation.

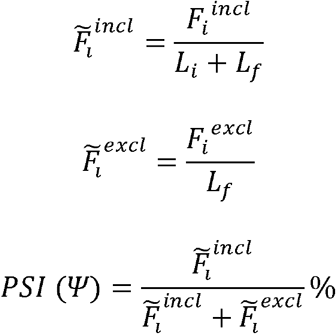

*F*_*i*_^*incl*^ incl number of reads or fragments supporting the inclusion of *i*-th exon; *F*_*i*_^*excel*^ number of reads or fragments supporting the exclusion of *i*-th exon; *L*_*f*_ fragment length; *L*_*i*_ size of *i*-th exon. We used PSI cutoffs of 20% and 80% to determine skipping and inclusion of exons based on the overall PSI distribution (Supplementary Figure 10).

### RNA-binding proteins

RBP enrichment was calculated based on the peaks identified from the eCLIP experiments. We downloaded the ENCODE eCLIP uniformly processed peaks from K562 and HepG2 cell types (see Supplementary Table 1 for eCLIP data accession). The peak was called using CLIPPER software [65] and filtered for peaks having a score of 1,000. We then counted numbers of RBP binding events at a base-pair resolution, agnostic to cell type.

To examine preferential binding patterns of splicing factors around SSs, RBP peaks were annotated as splicing-related factors if they belong to hnRNP- and SR-families (n=29). We extended both 3’ acceptor and 5’ donor SS by 1,000 bp in both up and downstream direction and binned the region into 100 bp intervals. We defined the position relative to the distance to the SS, in the 5’ to 3’ direction. For each interval, we calculated the frequency of splicing factor binding normalized to the size of the interval. The value of RBP enrichment means the normalized binding frequency of splicing-related factors.

### LSTM model

We adopted the following equations for the modeling of splicing using LSTM. σ function denotes sigmoid function. ⊗ denotes Hadamard product where two matrices are multiplied in a pair-wise fashion. *x*_*t*_ denotes input vector and *h*_*t*_ denotes output vector, *f*_*t*_ denotes forget gate vector, *i*_*t*_ denotes input or update gate vector, *o*_*t*_ denotes output gate vector, *c_t_* denotes cell state vector.

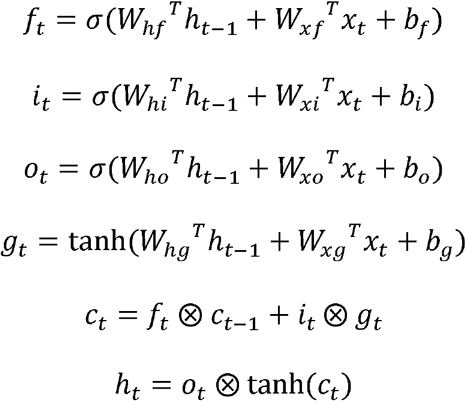

### GRU model

We adopted the following equations for the modeling of splicing using GRU. *x*_*t*_ denotes input vector and *h*_*t*_ denotes output vector, *z*_*t*_ denotes update gate vector and *r_t_* denotes reset gate vector.

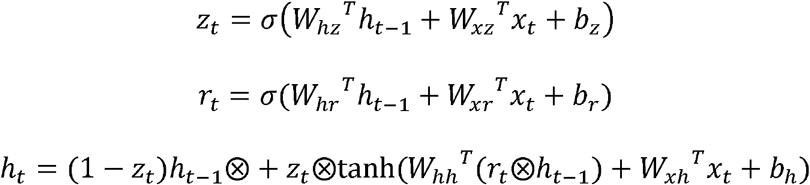

### Pre-processing of data for the training model

We selected six normal and three cancer samples from the reference epigenome series. The dataset contains consolidated epigenomes from the Roadmap Epigenomics Consortium [41] and the ENCODE Consortium. All datasets were uniformly processed and mapped to the GRCh38 human reference genome. All samples contained a core set of histone modification tracks (H3K4me1, H3K4me3, H3K36me3, H3K27ac, H3K27me3, and H3K9me3) as well as RNA-seq data. We used additional histone modification tracks, as well as DNase I hypersensitivity, DNA methylation, and nucleosome positioning tracks, to predict alternative splicing upon availability. Detailed information on datasets used can be found in Supplementary Table 1. For each exon, we obtained DNA sequences at intron-exon boundaries (3’ acceptors) and exon-intron boundaries (5’ donors), as well as 100 bp upstream and downstream of SSs. Splice junctions included both canonical and non-canonical SSs. We processed all sequences to read in the 5’ to 3’ direction using strand information from each gene. Each 400 bp DNA sequence was encoded into a 1,000 by 4 binary array using one-hot encoding. We used RNA-seq expression profiles to indicate tissue-specific alternative splicing patterns. Genes having fewer than two exons were discarded and the first and last exons were excluded from the analysis. We classified an exon as being expressed if its FPKM was greater than or equal to 1. We normalized all ChIP-seq histone modification tracks and DNase-seq tracks over corresponding input signal tracks using MACS v2.0.10 (https://github.com/taoliu/MACS) [66]. We used negative log10 of the Poisson p-value to measure the enrichment level over the background. Due to the wide dynamic range observed, we used a p-value threshold of 1e-2 for the upper limit. We processed all feature tracks including DNA methylation and nucleosome signal tracks to read in the 5’ to 3’ direction and scaled them to a range of 0 to 1.

### Performance evaluation of the model

There is no single metric that can give you a measure of performance in a binary classification problem. Relying on one metric can be misleading especially when there is high class imbalance. Therefore, we employed various metrics to measure the performance of the predictive model. ROC curve explains the tradeoff between true-positive rate (TPR) and false-positive rate (FPR). PR curve visualizes the tradeoff between positive predictive value (PPV) and true-positive rate (TPR).

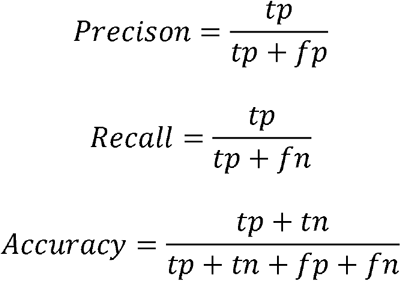

In addition, we used F1-score, which is the harmonic mean of precision and recall, to measure the performance of the splicing model.

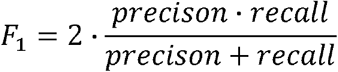

### Hyperparameter tuning of splicing model and training

We tested a range of dimensions and depths of RNN models and network design hyperparameters to optimize the alternative splicing model. We chose optimal hyperparameters by tuning one parameter at a time while fixing the rest. Hyperparameters included but were not limited to the number of recurrent layers, size of neurons in each layer, pooling strategy, dropout rate, choice of activation function and loss function, optimizer, and number of the epoch. We shuffled the order of the data and split the dataset into training and test sets using an 80 to 20% ratio. 20% of test data was set aside for the performance evaluation. 80% of training data was split again between 80 to 20% (64 and 16% of the original data) for fitting the model and validating the model fit during the training phase. We fed a range of sequences from 50 to 1,000 bp within each SS and found the 400 bp span to be the ideal size for the model. For the neural network architecture, we achieved the best result when two RNN units were stacked together, which allowed the model to learn higher-level temporal representations. We used a hidden state size of two by default and we recommend not using a hidden state size greater than 128 to avoid overfitting problems (Supplementary Figure 8A). We applied three variants of the RNN model, LSTM [45], GRU [67], and simple RNN. To compare the performance of memory-based units (LSTM and GRU), we implemented a simple RNN model using the same network architecture. We found that both LSTM and GRU were capable of learning long-term dependencies and were effective in learning high-dimensional contextual relationships between epigenomic features around the SSs. We split the input sequences into two parts where the first half represented a 3’ acceptor SS and the latter half represented a 5’ donor SS. We fed these sequences into two separate RNN units of size 200 and merged them into another RNN unit of size 400. The last RNN layer was followed by a dropout layer to prevent overfitting of the training dataset. The last fully-connected layer contained the softmax activation function for classifying exons as either spliced or unspliced. To train the model, we used a binary cross-entropy objective function with the Adam optimizer [68]. For each dataset, we trained the model for 20 epochs. We tested the implementation of ESPRNN using TensorFlow v2.0 (https://www.tensorflow.org). Our implementation also works with Keras v1.0.3 or v2.2.4 (https://github.com/fchollet/keras) with either TensorFlow v1.15 and Theano v0.8.2 [69] backend with a minor tweak. We used various Nvidia GPUs (Titan K20m, K80, GTX 1080ti, RTX2080, P100, and Titan V) to train the model.

## Acknowledgements

We would like to acknowledge Steve Weston from the Yale Center for Research Computing for technical support in setting up our GPU computing infrastructure.

## Supporting Information Legend

**Supplementary Table 1**

**List of datasets and the accession numbers used for the study.**

**Supplementary Table 2**

**Overview of dataset used for training the ESPRNN model.** The model was trained using the CORE (highlighted in red) and FULL set based on the availability of data. The CORE set was used to compare the predictive performance across cell types.

**Supplementary Table 3**

**ESPRNN model prediction performance measured by F1 score.** Predictive performance was compared between the CORE and FULL set of genomic features. For each set, performance was compared using LSTM, GRU, and simple RNN models. Predictive performance was measured by F1 score.

**Supplementary Table 4**

**Comparison of models trained with 50 bp span and 100 bp span data.** Each model was trained using genomic features derived from 50 bp span or 100 bp span data from splice sites using the LSTM model. Performance was measured using F1 score and ROC AUC.

**Supplementary Figure 1.**
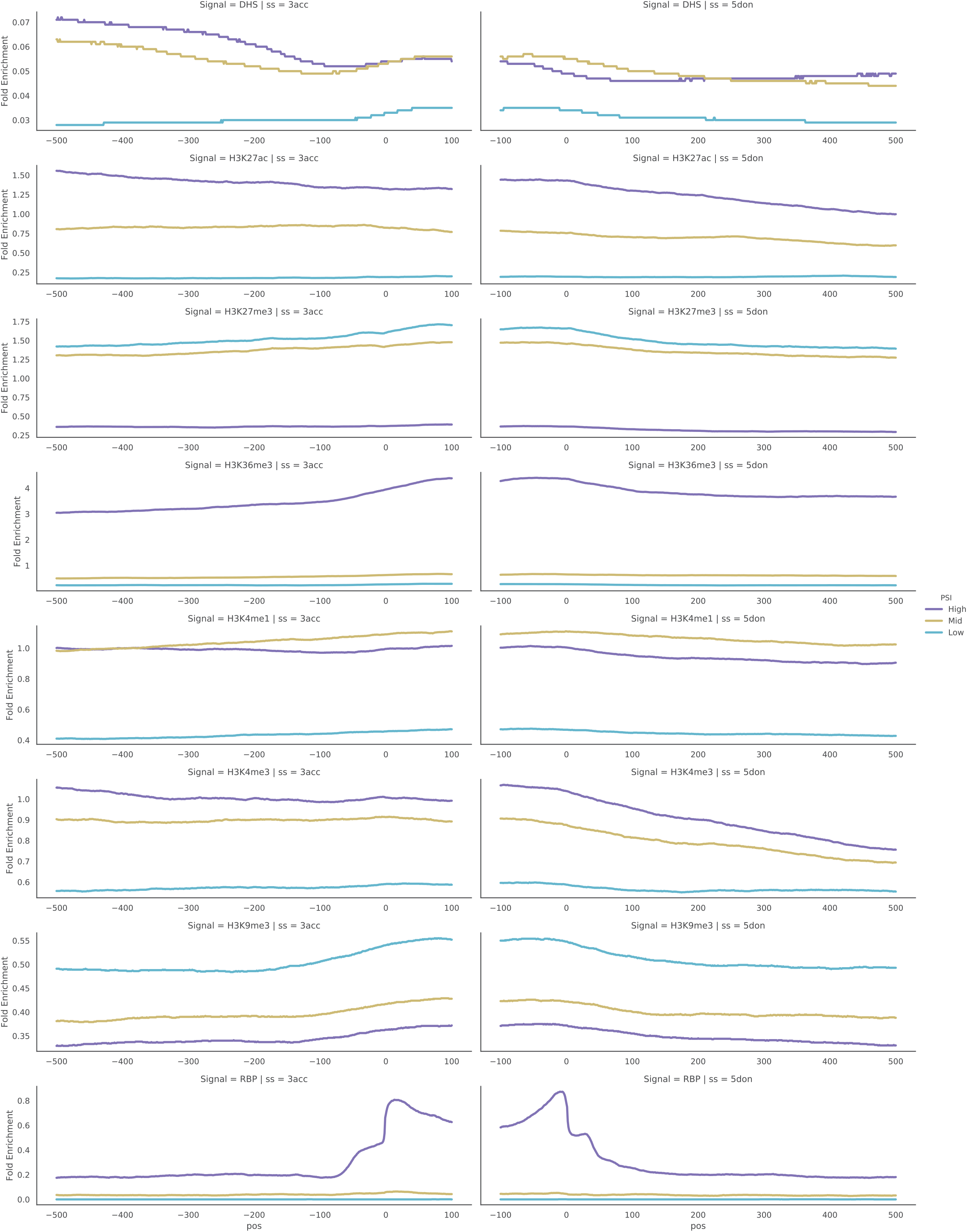
(Shadow figure of the main Figure 2A) Enrichment of various epigenomic marks of HepG2 at the exon-intron boundary. High PSI indicates exon inclusion, mid PSI indicates exons with 40-60% PSI, and low PSI indicates exon skipping.

**Supplementary Figure 2.**
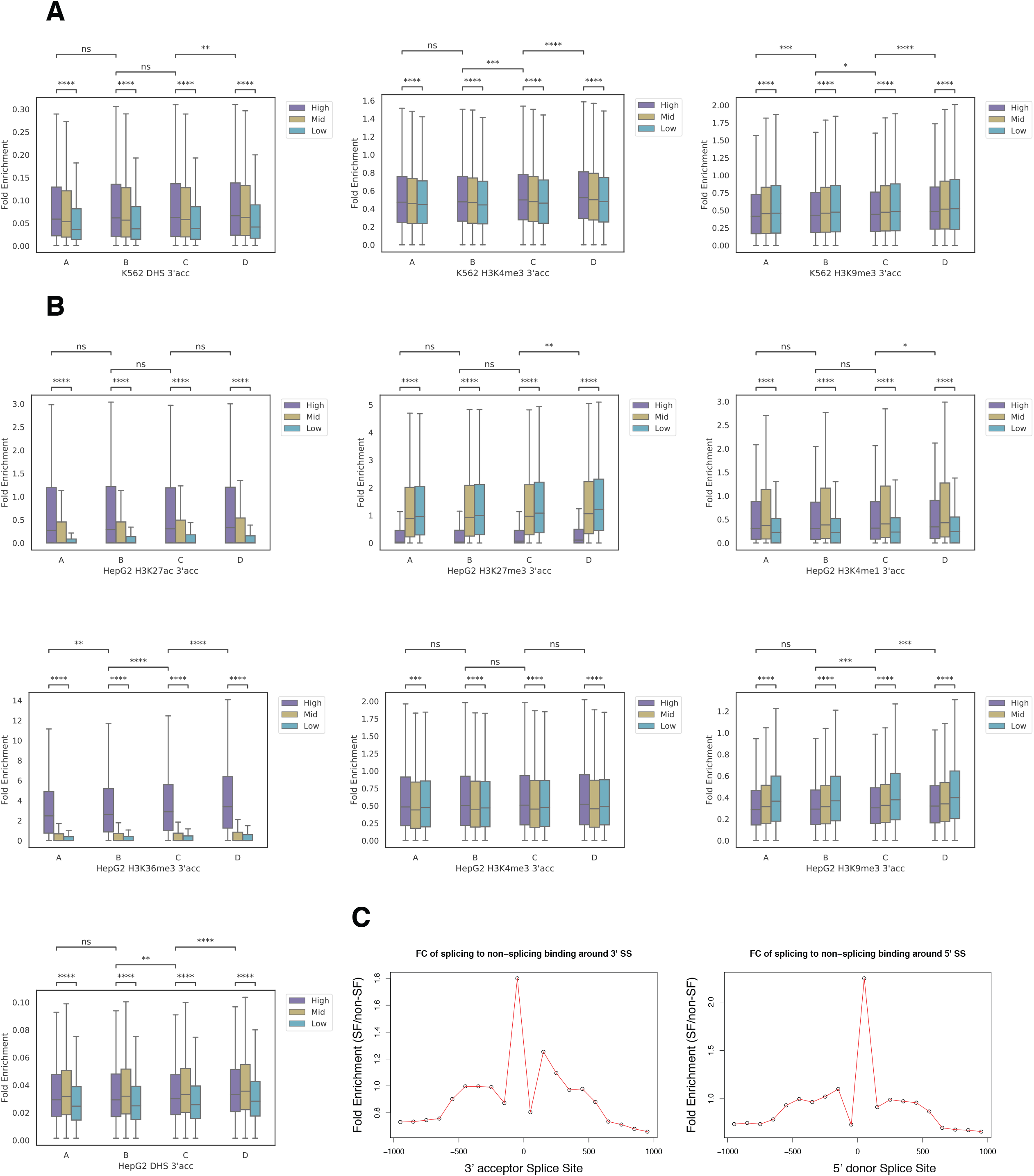
(Shadow figure of the main Figure 2B) Comparison of epigenetic enrichment around different segments of the 3’ acceptor site for **(A)** K562 and **(B)** HepG2. High PSI indicates exon inclusion, mid PSI indicates exons with 40-60% PSI, and low PSI indicates exon skipping. Mann-Whitney-Wilcoxon two-sided test, ns: 0.05 < p <= 1; *: 0.01 < p <= 0.05; **: 0.001 < p <= 0.01; ***: 0.0001 < p <= 0.001; ****: p <= 0.0001. **(C)** Fold enrichment of splicing-related RBPs to non-splicing-related RBPs around the 3’ acceptor splice site and 5’ donor splice site.

**Supplementary Figure 3.**
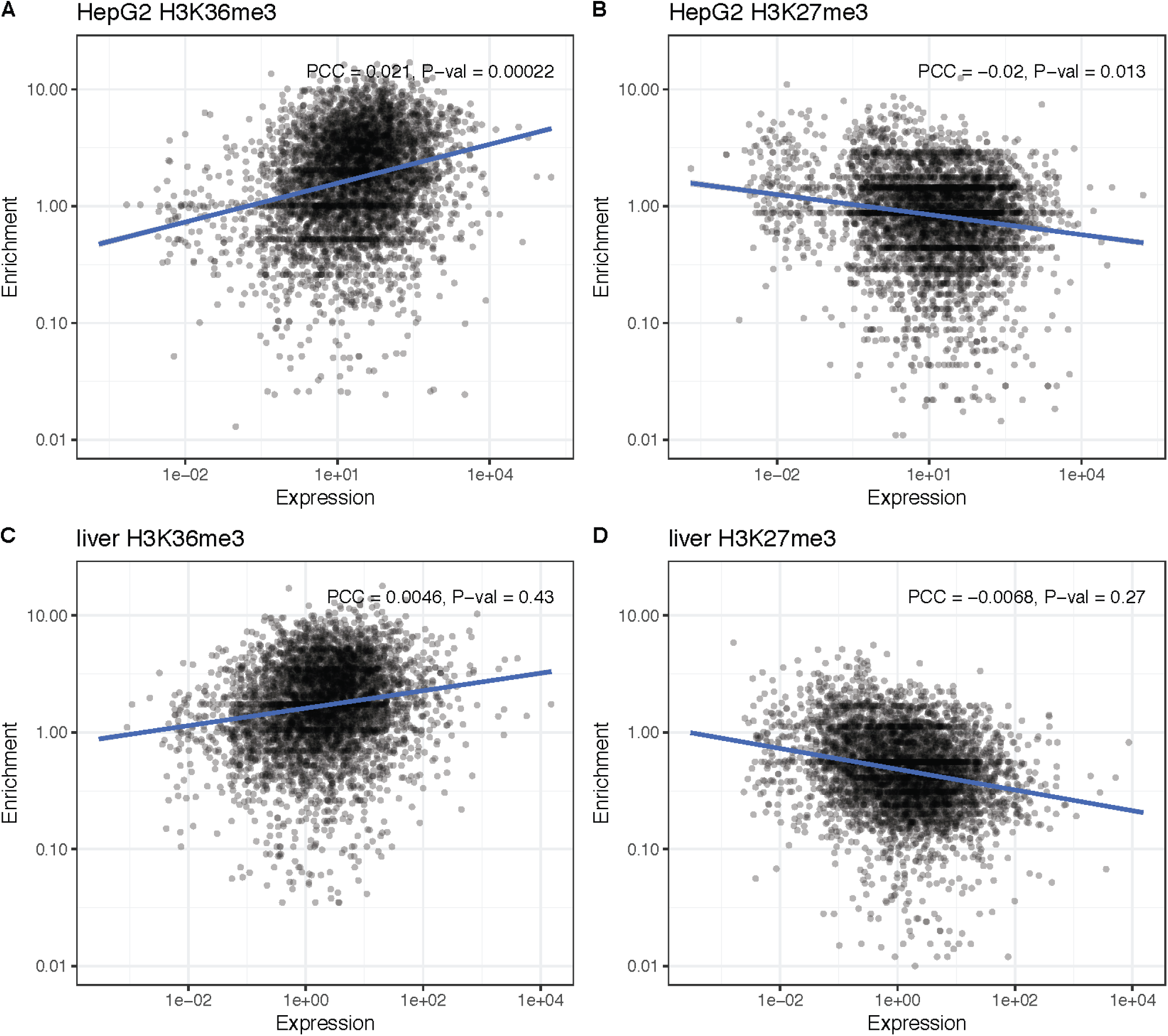
Correlation of exonic expression (FPKM) and histone enrichment of **(A)** HepG2 H3K36me3, **(B)** HepG2 H3K27me3, **(C)** liver H3K36me3, and **(D)** liver H3K27me3. PCC: Pearson Correlation Coefficient.

**Supplementary Figure 4.**
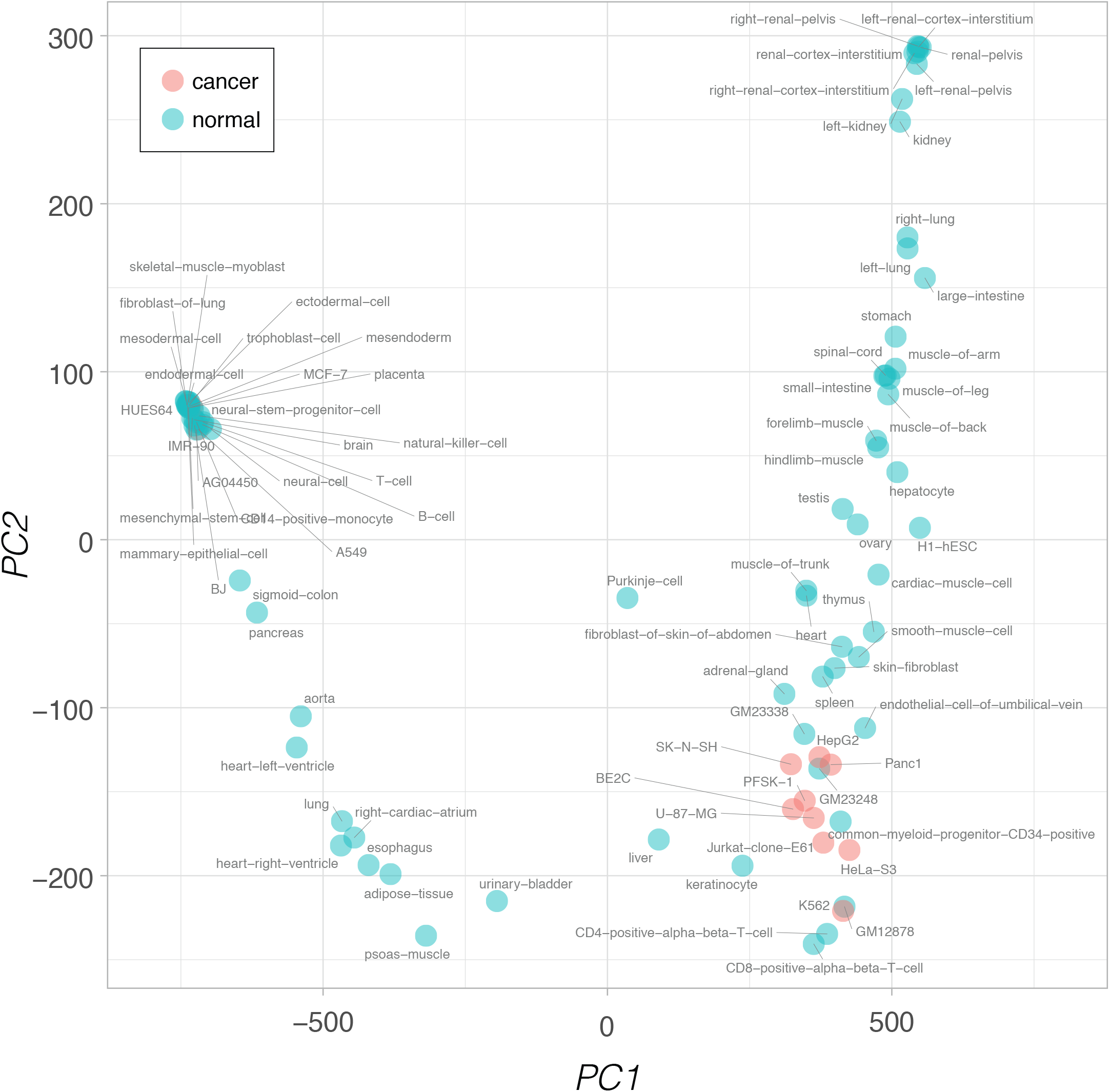
Splicing patterns based on exonic expression level (FPKM) for diverse ENCODE cell types are projected on a PCA cell space.

**Supplementary Figure 5.**
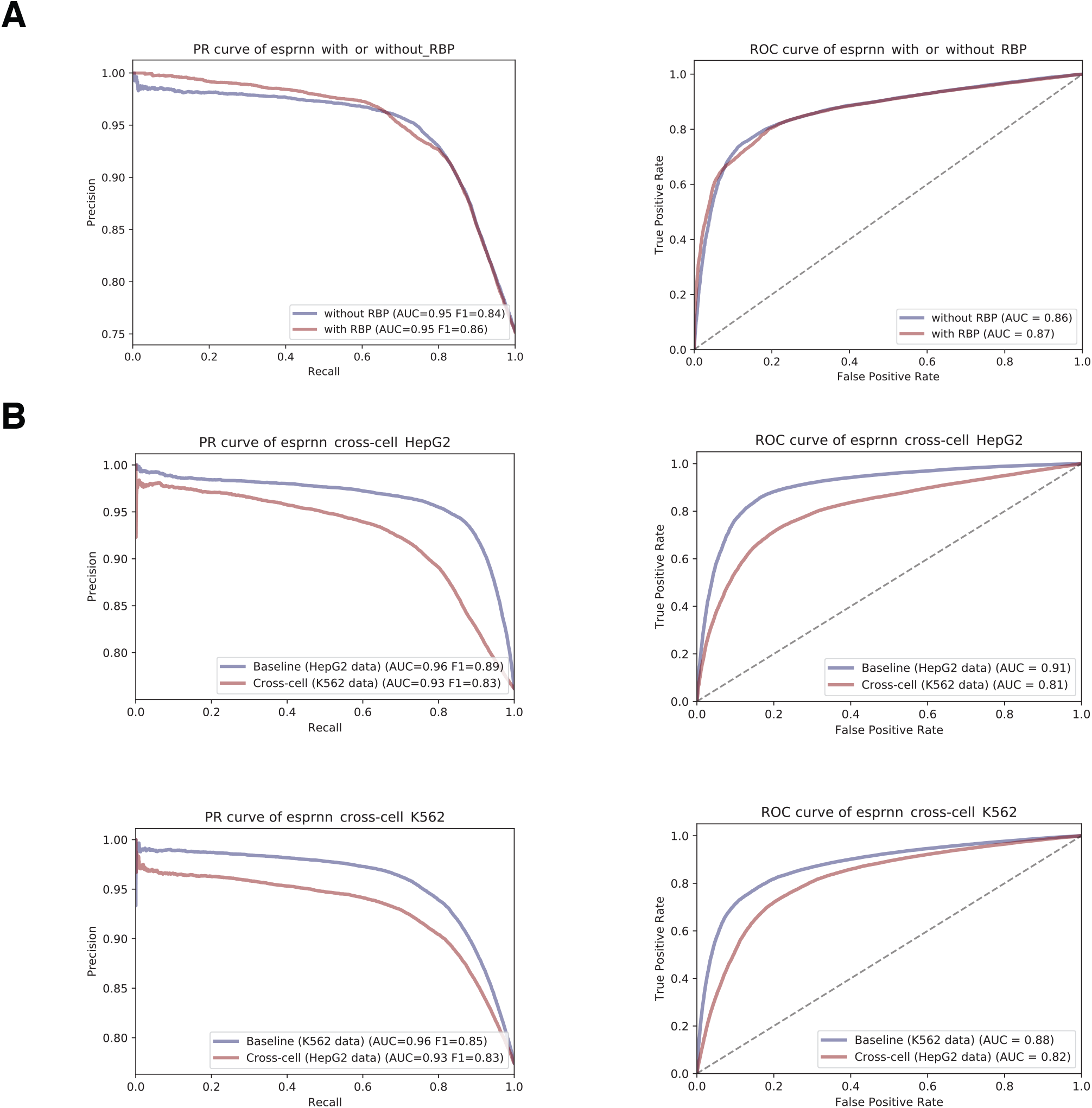
**(A)** Difference in splicing prediction performance when RBP binding profiles were added as an additional feature of the base model containing chromatin accessibility and histone marks. **(B)** Cross-cell testing of model. Model was trained on HepG2 data and tested on K562 data, and vice versa.

**Supplementary Figure 6.**
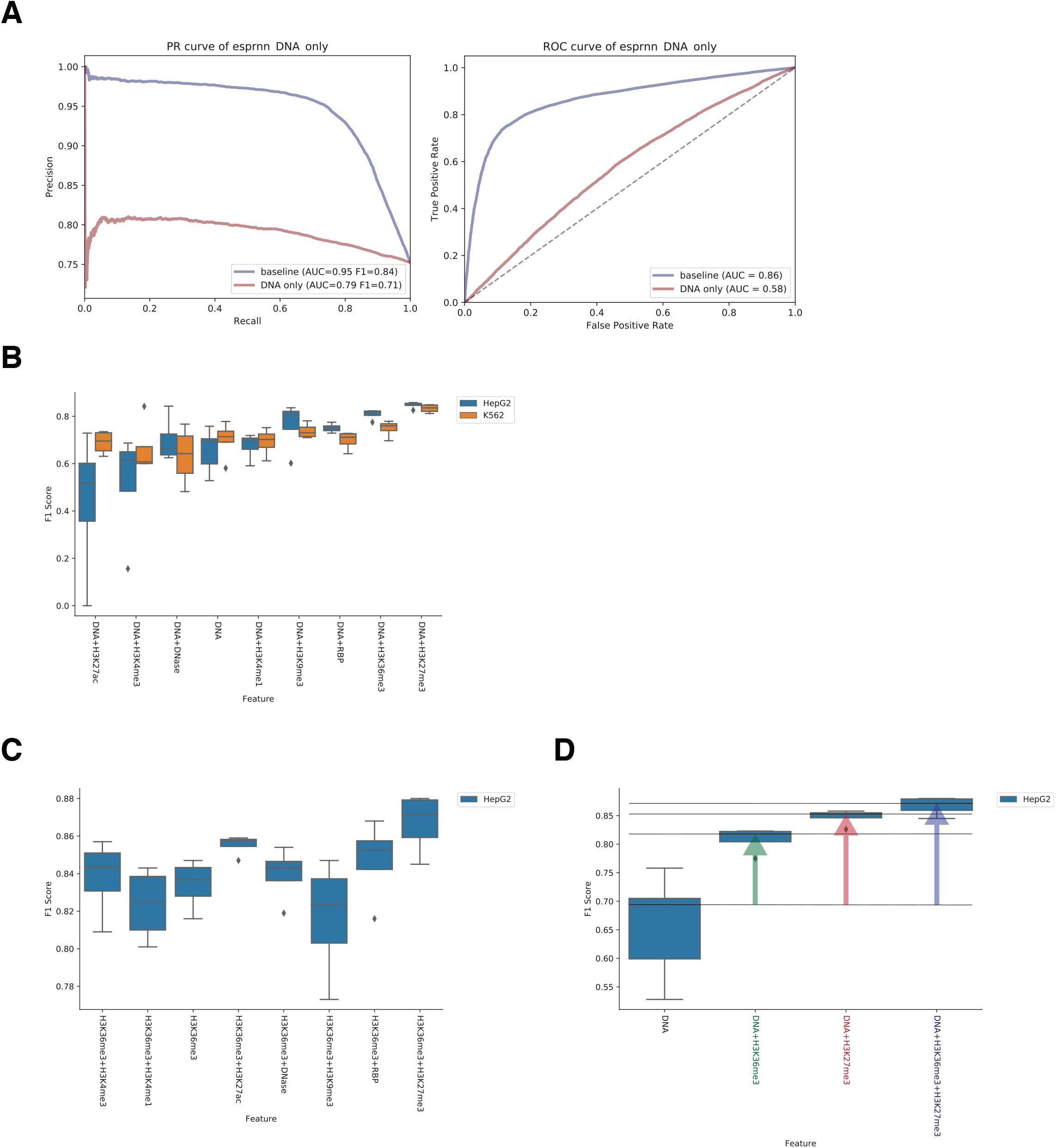
**(A)** Comparison of the baseline model trained using chromatin accessibility and 6 histone marks to a model using DNA sequence feature only **(B)** Measure of information gain from additional epigenetic feature based on DNA sequence only model **(C)** Comparison of splicing prediction performance using a pair of epigenetic features. (D) Performance comparison of models using H3K36me3 or H3K27ac feature individually to a model using both H3K36me3 and H3K27ac features. Performance was measured based on F1 score from 5 trials.

**Supplementary Figure 7.**
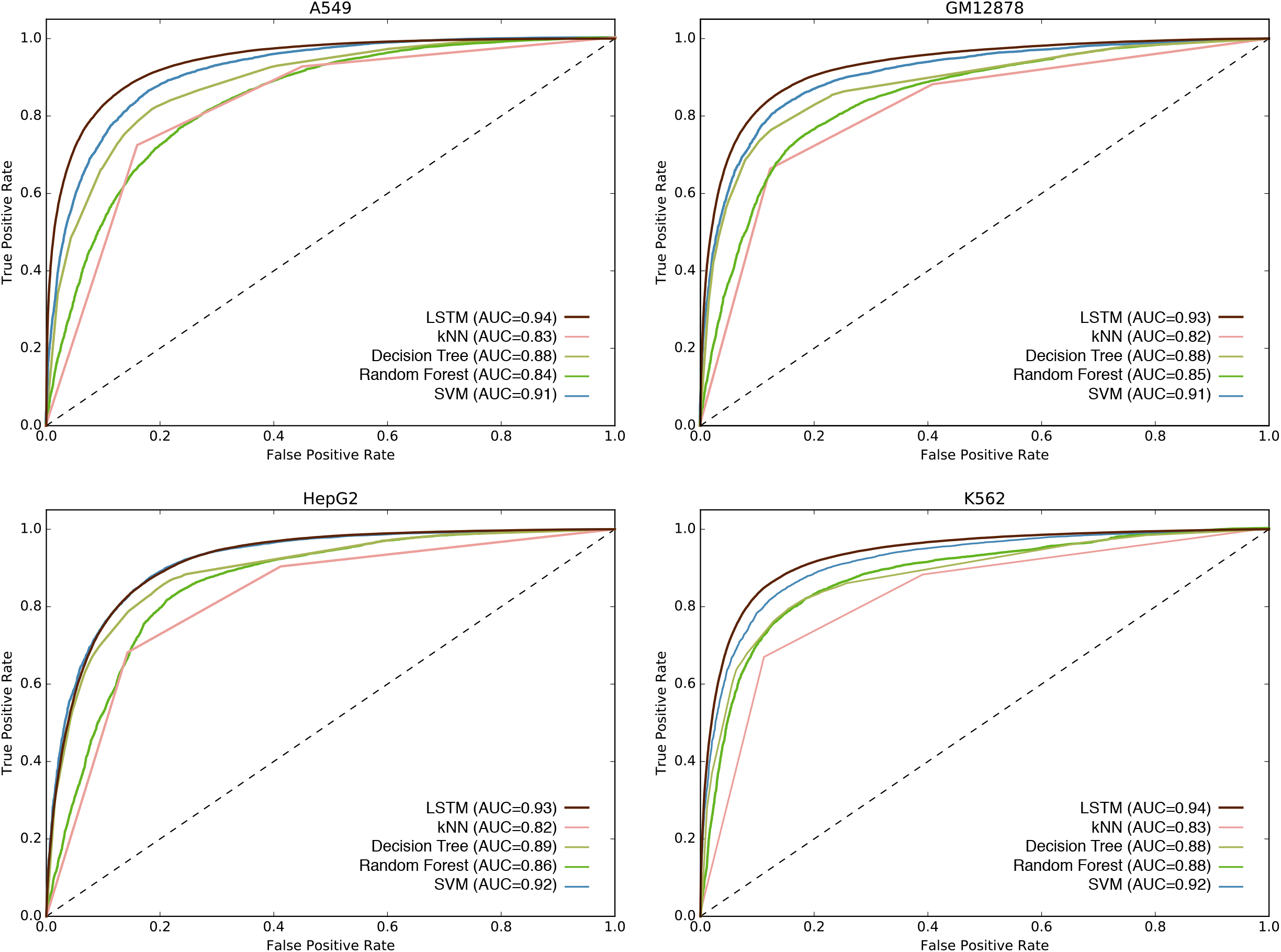
Comparison of LSTM-based model with other machine learning algorithms. Four different algorithms, k-Nearest neighbor (kNN), decision tree, random forest, and support vector machine (SVM), were compared to the LSTM-based model across four different tissue types (A549, HepG2, GM12878, K562).

**Supplementary Figure 8.**
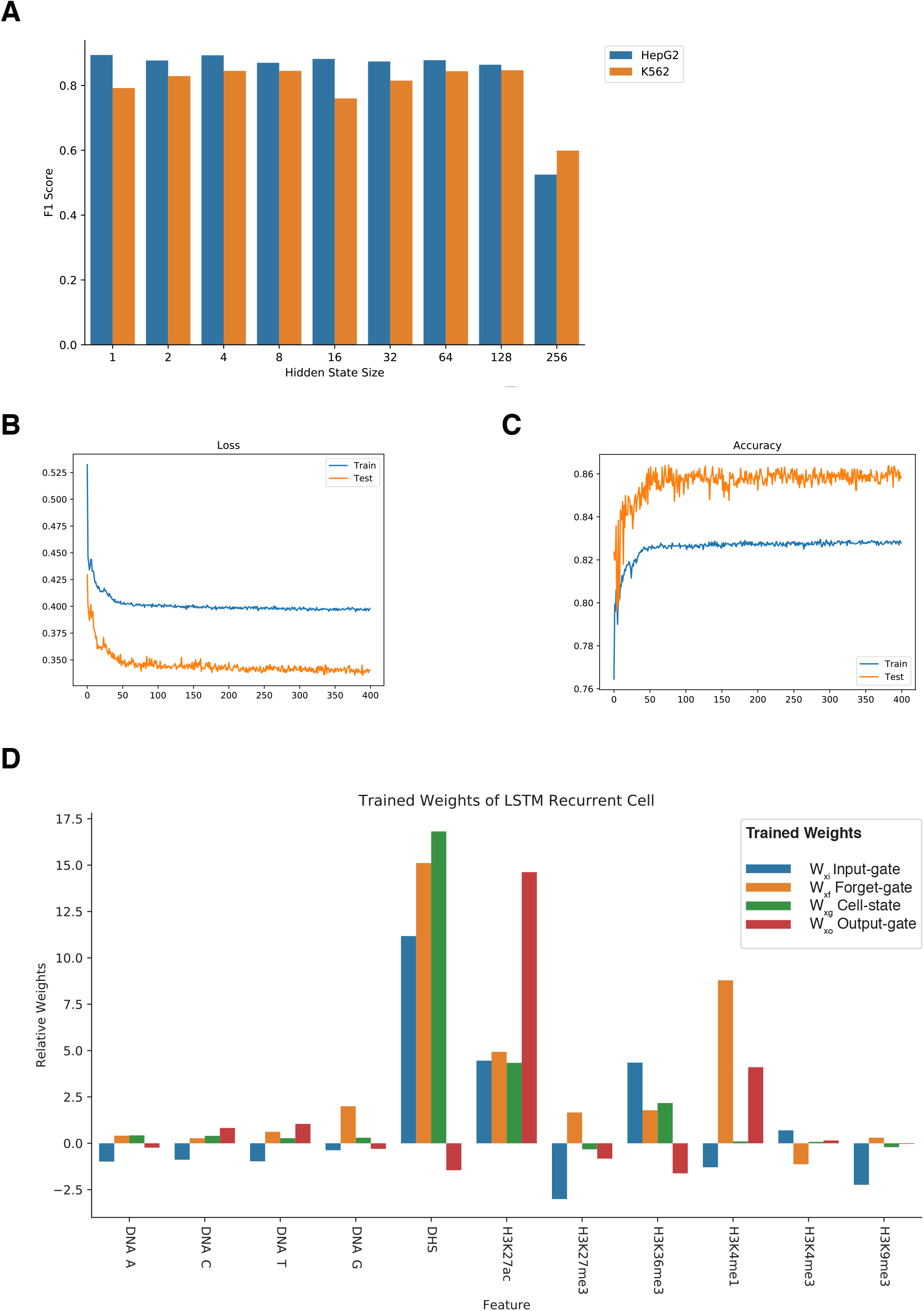
**(A)** Comparison of splicing prediction performance across different sizes of hidden state. **(B)** Loss of training an LSTM model with 1 hidden layer for 400 epochs. **(C)** Accuracy of training an LSTM model with one hidden layer for 400 epochs. **(D)** Trained weights of LSTM recurrent cells.

**Supplementary Figure 9.**
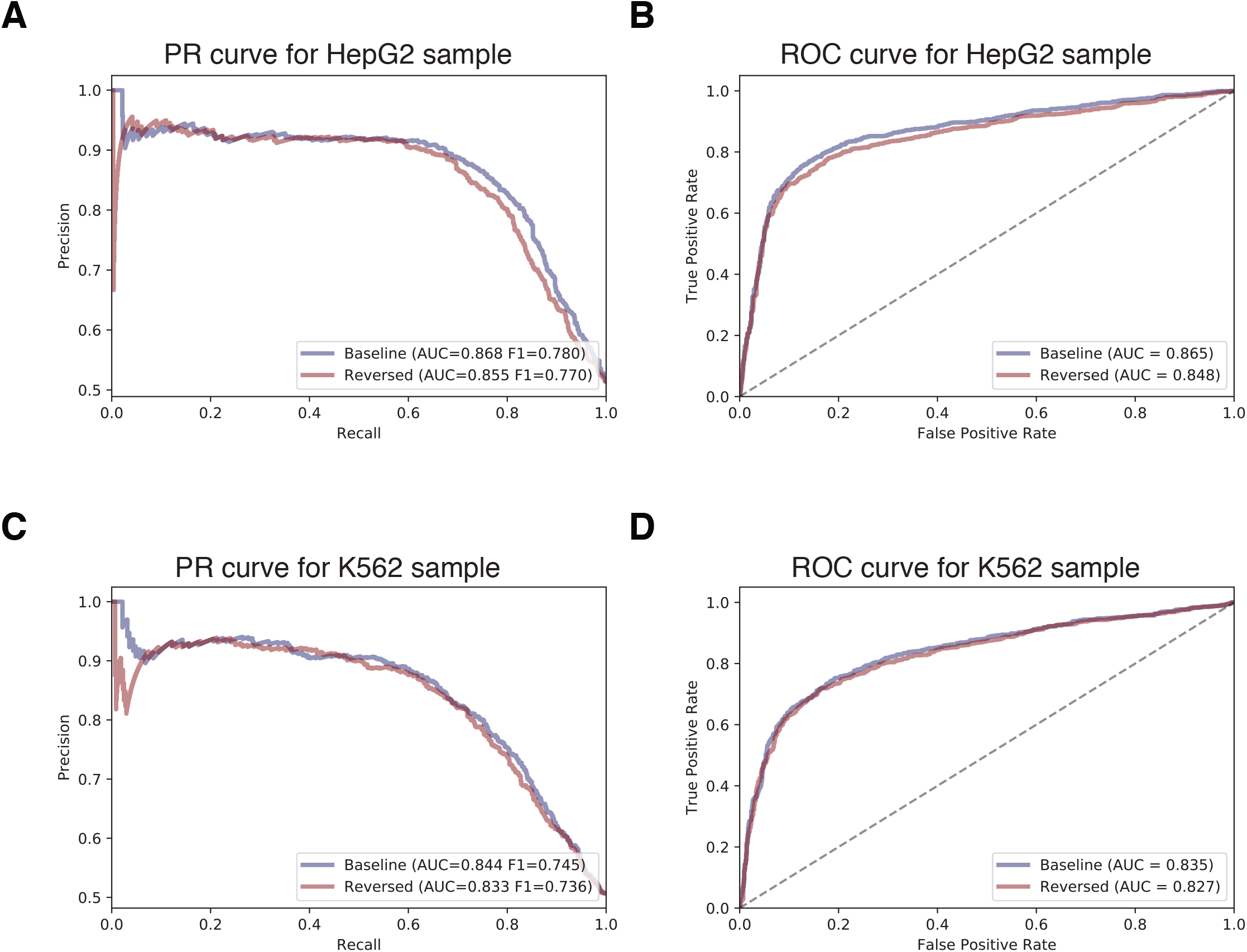
Comparison of splicing prediction performance when epigenetic context features are reversed in time-direction. **(A)** precision-recall curve for HepG2 **(B)** ROC curve for HepG2 **(C)** precision-recall curve for K562 **(D)** ROC curve for K562

**Supplementary Figure 10.**
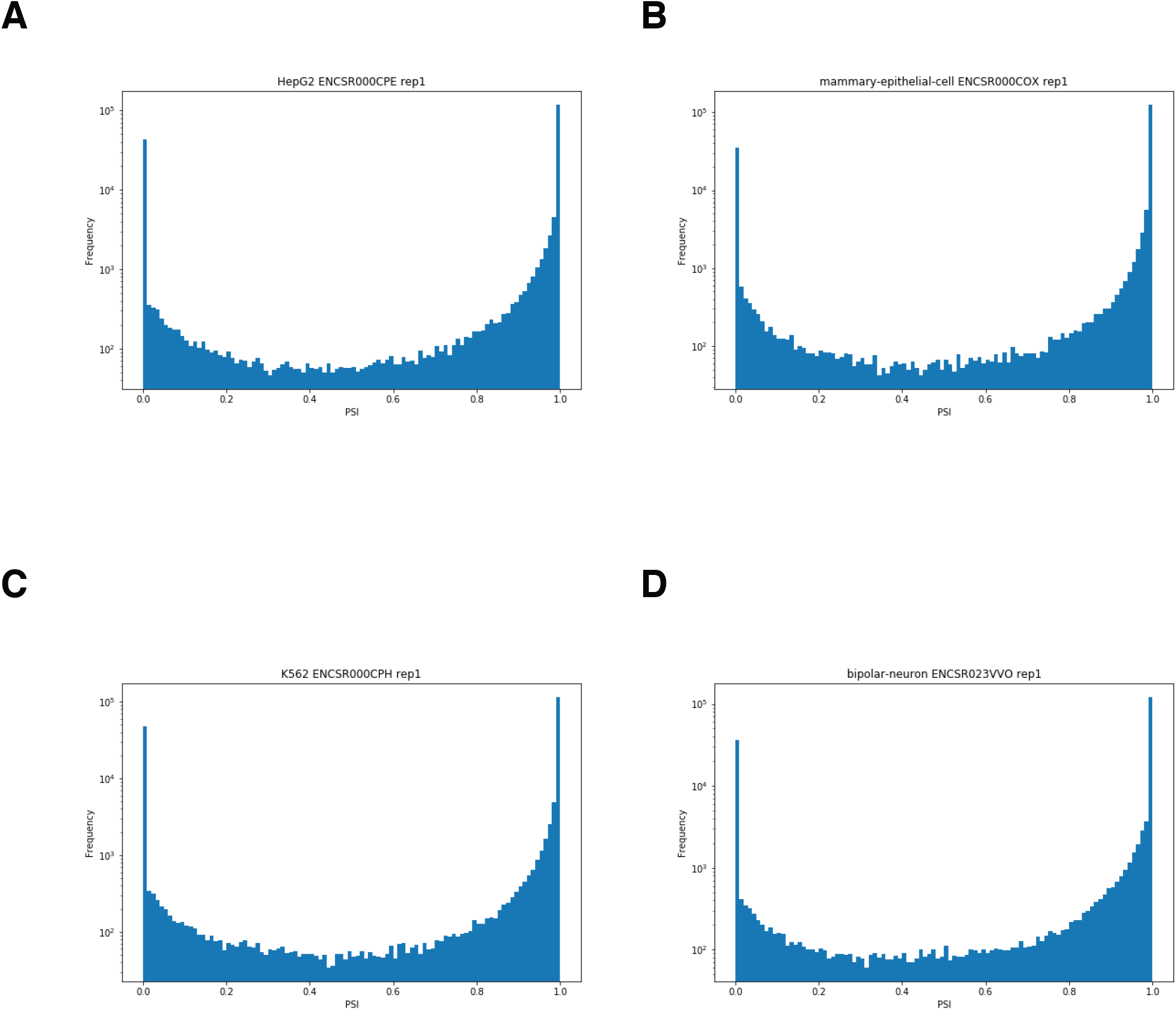
PSI histogram of cassette exons from **(A)** HepG2 **(B)** mammary epithelial cell **(C)** K562, and **(D)** bipolar neuron.

